# RNase E biomolecular condensates stimulate PNPase activity

**DOI:** 10.1101/2022.06.06.495039

**Authors:** Michael Collins, Dylan T. Tomares, Jared M. Schrader, W. Seth Childers

## Abstract

Bacterial Ribonucleoprotein bodies (BR-bodies) play an essential role in organizing RNA degradation via liquid-liquid phase separation in the cytoplasm of bacteria. BR-bodies mediate multi-step mRNA decay through the concerted activity of the endoribonuclease RNase E coupled with the 3’-5’ exonuclease Polynucleotide Phosphorylase (PNPase). Our past *in vivo* studies indicated that the loss of PNPase recruitment into BR-bodies led to a significant build-up of RNA decay intermediates in *Caulobacter crescentus*. We reconstituted RNase E’s C-terminal domain together with PNPase to understand how RNase E biomolecular condensates can tailor the functions of PNPase. We found that PNPase catalytic activity is accelerated when colocalized with the RNase E biomolecular condensates. In contrast, disruption of the RNase E-PNPase protein-protein interaction led to a loss of PNPase recruitment into the BR-bodies and a loss of ribonuclease rate enhancement. We also found that BR-bodies could enhance the decay of select RNA substrates, as we observed a 3.4-fold enhancement of polyadenylic acid (poly(A)) degradation and no impact upon poly(U) degradation. Our investigation into the origins of the 3.4-fold rate enhancement for poly(A) decay indicates a combination of scaffolding and mass action effects impact due to the concentrated biomolecular condensate environment accelerating RNA decay. Consistent with our past *in vivo* work, these studies suggest BR-bodies are sites of accelerated RNA decay that can shape the available transcriptome.

## Introduction

Biomolecular condensates are liquid-like to gel-like protein assemblies that form membraneless compartments that organize multi-step biochemical pathways within cells^1^. Within these protein-rich ensembles, scaffolding proteins exhibit weak multivalent protein-protein^2^ and protein-nucleic acid interactions^3^ that facilitate phase separation into liquid-like droplet assemblies. The scaffold also recruits client proteins into these assemblies resulting in functional compartments that include stress granules and p-bodies^4, 5^, the nucleolus^4, 6^ and signaling complexes^6, 7^. Recently, bimolecular condensates have been discovered as a way in which bacteria organize biochemistry into organelle-like structures^8, 9^. This has opened the door to reconsidering microbial biochemistry in the context of non-membrane bound compartments.

Many bacterial biomolecular condensates have been discovered in diverse biochemical pathways^8^. These membraneless compartments organize ABC transporters^10^, single-stranded (ss) DNA-binding proteins^11^, aggresomes^12^, cell division protein FtsZ^13^, BapA amyloid biofilm matrix protein^14^, circadian rhythm associated proteins^15^ and carboxysomes^16^. For example, in *Caulobacter crescentus*, two compositionally distinct biomolecular condensates regulate a network of signaling proteins to promote asymmetric cell division^17, 18, 19^. It has been shown that histidine kinases can be regulated spatially by sensory domain stimulation^17^. Moreover, low-phosphate nutrient conditions modulate levels of ATP, which directly impacts the phase separation properties of the scaffold that regulates histidine kinase activity^19^. Much like the intersection of nucleic acids and biomolecular condensates in mammals, the earliest discovered bacterial biomolecular condensates involved protein scaffolds that sequester nucleic acids as clients^20-22^. For example, our past studies have discovered that the *C. crescentus* degradosome phase separates as bacterial ribonucleoprotein bodies (BR-bodies) that mediate the rapid decay of RNAs^20, 21^.

Here we investigated how Ribonuclease E (RNase E) regulates the enzymatic functions of PNPase^20, 21^. RNase E and PNPase are two major proteins that make up the RNA degradosome, an interaction broadly conserved within many but not all bacterial species^23^. RNase E contains an N-terminal endoribonuclease domain and a C-terminal disordered domain that scaffolds RNAs, PNPase, RNase D, and aconitase (Figure 1)^24^. Studies have successfully reconstituted the RNA degradosome complex in *E. coli*^25^. This includes seminal work by Mackie and colleagues that demonstrated complete degradation of the structured *malEF* substrate required the concerted functions of *E. coli* RNase E, RhlB, and PNPase^26^. Other studies have built upon this work showing the critical roles of RhlB, PNPase and RNase E working as a complex to degrade structured RNAs^27^. However, the role of phase separation on PNPase activity has not been studied *in vitro*.

**Figure 1:**
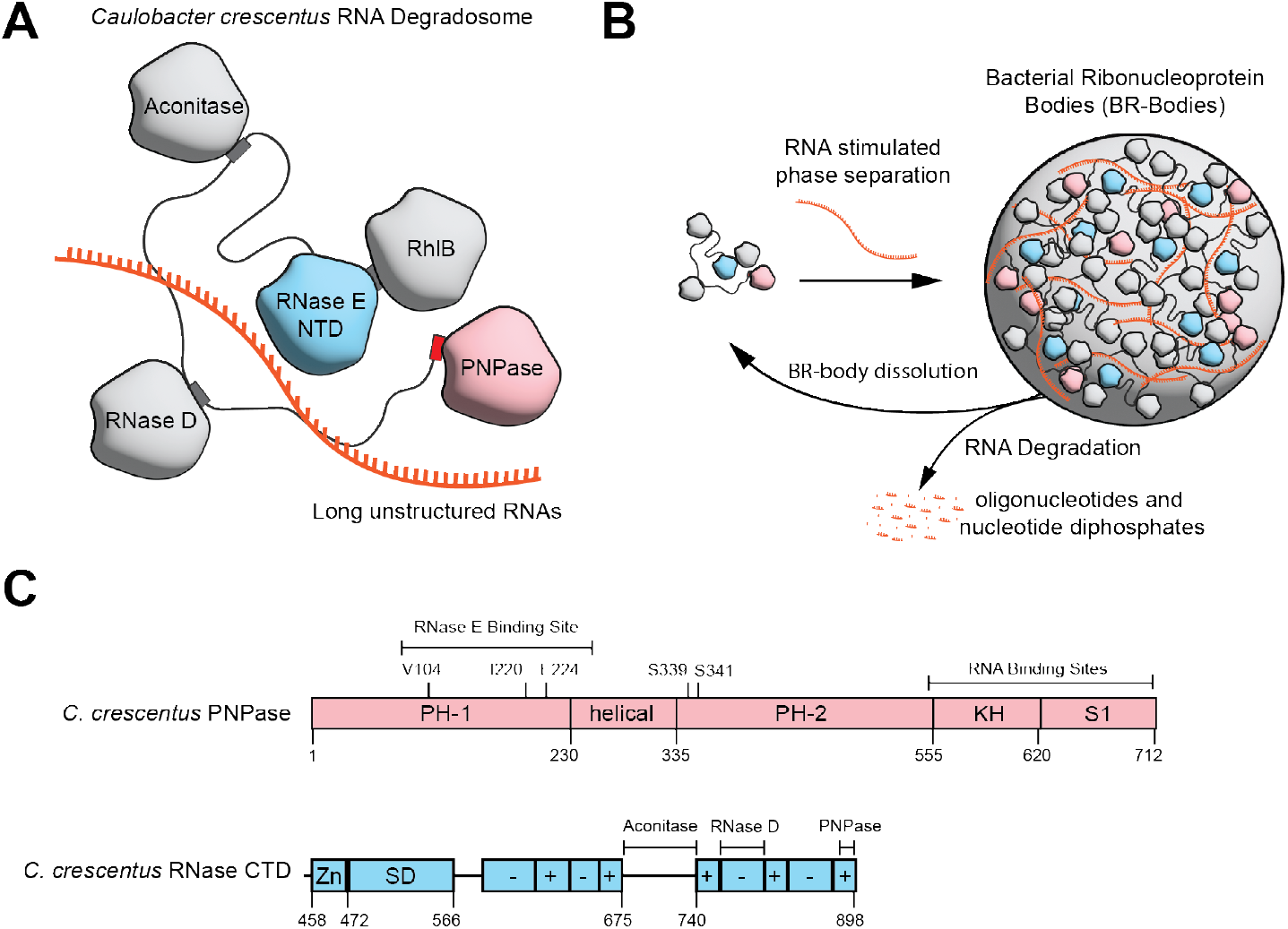
Bacterial Ribonucleoprotein Bodies (BR-bodies) are phase-separated biomolecular condensate of the RNA degradosome. (A) The RNA degradosome in *Caulobacter crescentus* consists of RNase E, which serves as the primary scaffold that recruits long unstructured RNAs, PNPase, RNase D, and Aconitase as clients. (B) *In vivo* phase separation of the degradosome is stimulated by multivalent interactions with the arginine-rich charge blocks^29^ in its C-terminal IDR and long unstructured RNA substrates resulting in the formation of BR-bodies. BR-bodies serve as sites of RNA degradation, in which the endoribonuclease RNase E performs the rate-limiting initial cut of the RNAs. Subsequently, PNPase exoribonuclease activity breakdowns the RNA intermediates. Breakdown of the RNAs into small oligoribonucleotides and nucleotide diphosphates results in a loss of multivalency and dissolution of the BR-bodies^34^ (C) Domain architecture of C. crescentus PNPase and RNase E. PNPase is composeD of RNase PH domains, the helical domain, the RNA-binding K-homology domain and S1 RNA-binding domains^30^. The RNase E C-terminal domain is sufficient for phase separation of BR-bodies.^29^ The RNase E CTD comprises a Zn-link, a small domain and an intrinsically disordered region organized as a set of charge blocks.

Our past studies showed that the *C. crescentus* RNase E phase separated and underwent liquid-like fusion events *in vivo* and *in vitro* (Figure 1) ^20^. Protein-rich droplets formed transiently in an RNA-dependent manner, as rifampicin-mediated inhibition of RNA polymerase reduced foci formation of RNase E *in vivo*^*20*^. *In vivo*, BR-body enrichment assays indicate that BR-bodies engage a broad set of RNA substrates with a preference for long and unstructured RNAs^21^. The role of RNase E in RNA degradation was highlighted by *in vivo* Rif-seq experiments, where studies indicated that failure to form biomolecular condensates and recruit exoribonuclease clients increased global RNA half-lives^21^. Specifically, a *C. crescentus* RNase E NTD-only variant, lacking phase separation capabilities and PNPase recruitment, showed a bulk increase in global RNA decay half-lives from 3.6 to 4.8 min^21^.

*In vivo* RNA decay profiling identified that degradosome protein recruitment involving PNPase was required to catalyze the second step of RNA decay, the rapid cleavage by PNPase^28^. Therefore, we investigated whether PNPase’s organization within BR-bodies stimulates this critical second step as the relative contributions of allostery, scaffolding, and phase separation are unknown. We provide direct *in vitro* evidence on how RNase E regulates the functions of PNPase.

## Results

### Recruitment of PNPase into the RNase E droplets requires the C-terminal binding site

To investigate the recruitment of PNPase into RNase E droplets, we purified a PNPase active site mutant (PNPase-S339A/S340A/S341A-mCherry) called PNP-ASM-mCherry. Here, we chose to purify the active site mutant to minimize the potential impact of PNPase’s breakdown of RNA upon co-localization within the RNase E biomolecular condensates. In addition, we purified the RNase E C-terminal domain (residues 451-898), which is sufficient for phase separation^29^, called RNase E-CTD, to consider how phase separation impacts PNPase activity. We visualized each mixture via phase contrast and fluorescence microscopy imaging. We then calculated the fluorescence intensity ratio in the concentrated versus the dilute phase for each assembly, which we term partitioning ratio (PR). Differences in the degree of protein enrichment in the dense phase impact the change in fluorescence intensity. However, the biomolecular condensate’s unique chemical environment may alter the refractive index or impact the quantum yield of fluorescent proteins. In addition, the increased fluorescent protein concentration may lead to the quenching of the fluorescence signal. Therefore, the PR reflects the combinations of these effects. We found 20 µM RNase E-eYFP in 10 mM MgCl_2_ and 100 mM NaCl phase-separated into protein-rich biomolecular condensates with a 3.6 ± 0.3 partitioning ratio (Figure 2A and 2B). The addition of the 5 µM PNPase-ASM client had no impact on the observed partitioning of RNase E (p > 0.05) (Figure 2B).

**Figure 2.**
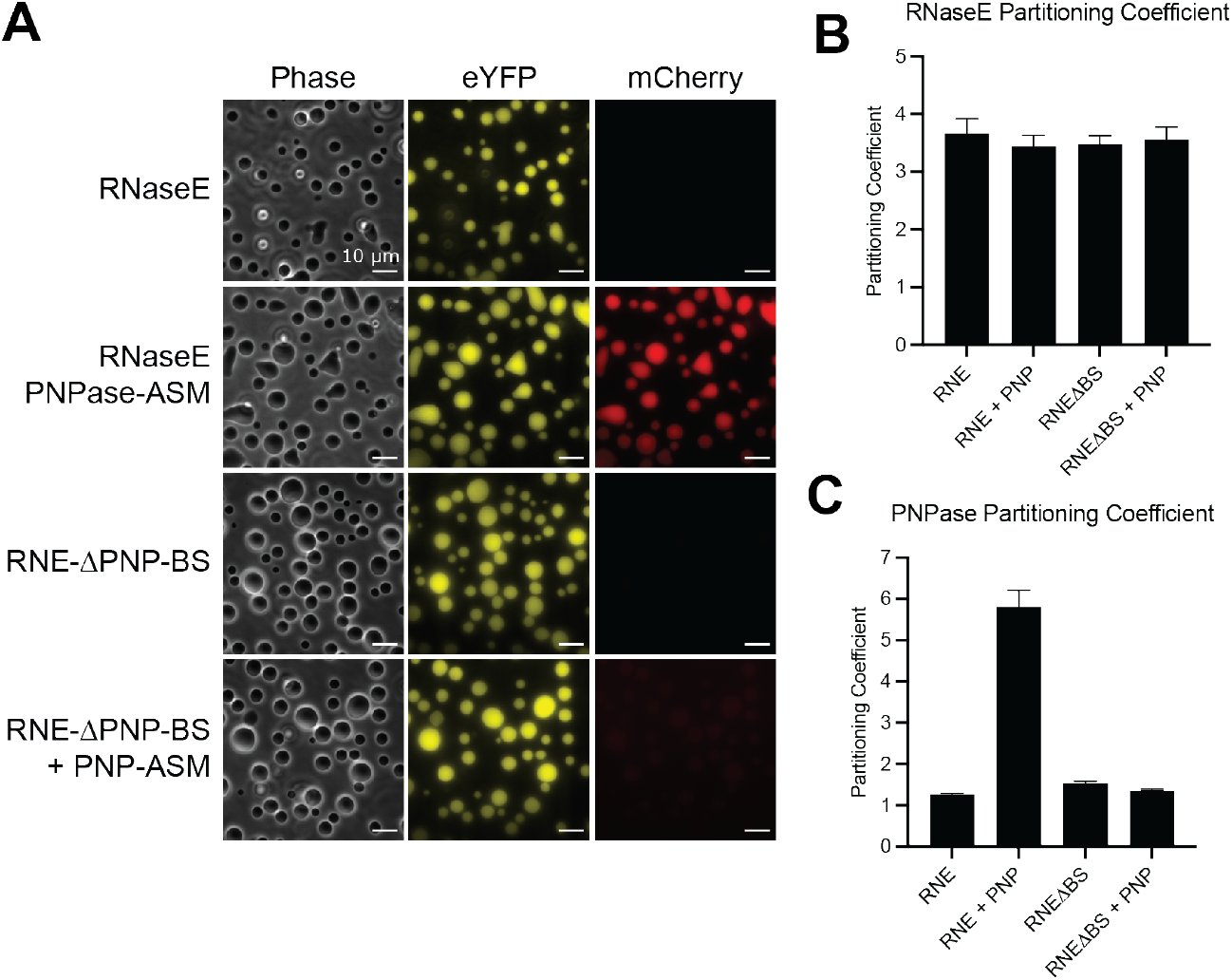
PNPase partitions into BR-bodies through an interaction with a C-terminal binding site on RNase E-CTD. (A) Phase contrast and fluorescence microscopy images of RNase E biomolecular condensates. RNase E and RNase E-ΔPNP-BS contain a C-terminal eYFP tag and were present at 20 µM, and PNPase-ASM (active site mutant) contains a C-terminal mCherry tag and was at a concentration of 1 µM. The scale bar is 10 µm. (B) Average partitioning ratios with their standard deviations are presented for RNase E. There is no statistically significant difference between any PRs (p > 0.05). (C) Average partitioning ratios with their standard deviations are presented for PNPase. Amongst experiments, PNPase is only significantly recruited into RNase E biomolecular condensates (p < 0.001). In all other cases, PNPase does not significantly partition into RNase E biomolecular condensates (p > 0.05). Data represent the average and standard deviation of partitioning ratios of n > 300 droplets.

PNPase-SM-mCherry by itself did not phase separate into a protein-rich phase under these conditions. However, the co-assembly of PNPase with RNase E, led to the recruitment of PNPase into the RNase E droplets with a partitioning ratio of 5.7 ± 0.7 (Figure 2C). We then tested if an RNase E variant, RNase E-ΔPNP-BS, lacking the 10 C-terminal residues that bind PNPase^30^, could recruit PNPase. Individually, both RNase E-CTD and RNase E-ΔPNP-BS phase-separated into biomolecular condensates (Figure 2A) with a partitioning ratio of 3.6 ± 0.3 and 3.5 ± 0.4, respectively. (Figure 2B). This indicates that the C-terminal PNPase binding site residues are unnecessary for RNase E’s homotypic phase separation. However, the RNase E-ΔPNP-BS variant displayed reduced capabilities to recruit PNPase with a partitioning coefficient of 1.4 ± 0.1, which was significantly less than RNase E-CTD recruitment of PNPase (p < 0.001) (Figure 2C). Thus, PNPase requires the 10 C-terminal residues of RNase E for enrichment within RNase E biomolecular condensates.

To understand the contributions of weak fluorescent protein interactions to recruitment within the RNase E biomolecular condensates, free mCherry was incubated with RNase E-CTD or RNase E-ΔPNP-BS. Free mCherry at 1 µM was poorly recruited into RNase E-CTD and RNase E-ΔPNP-BS with partitioning ratios of 1.2 ± 0.1 or 1.2 ± 0.1, respectively (Figure S1 A-C). This indicates that weak enrichment with partitioning ratios in the range of 1.0-1.3 may be due to weak fluorescent protein interactions. In contrast, enhanced recruitment beyond that amount requires a specific binding site.

### PNPase triple mutant disrupts recruitment into RNase E biomolecular condensates

Deleting the C-terminal residues of RNase E may alter the multivalent contacts that mediate RNase E phase separation. Therefore, we considered if mutations within PNPase could disrupt recruitment into the wild-type RNase E-CTD biomolecular condensates. RNase E binds to a hydrophobic pocket on the external surface of the catalytic core of PNPase facilitated by residues G896, W897, and W898^30^. These RNase E residues interact directly with PNPase’s V104, I220, E224, and F233 residues (Figure 3A). We hypothesized that mutating these PNPase residues would diminish PNPase recruitment into the RNase E biomolecular condensates. Therefore, we cloned and purified a PNPase variant, PNPase-V104A/E224A/F233A, to disrupt interactions between RNase E and PNPase. When incubated with RNase E, incorporation of PNPase-V104A/E224A/F233A into the BR-bodies was diminished (Figure 3B) from a partitioning coefficient of 4.4 ± 0.2 (PNPase-ASM) to 1.0 ± 0.1 (PNPase-V104A/E224A/F233A) (Figure 3C). Thus, PNPase utilizes a specific binding site for enrichment into RNase E biomolecular condensates.

**Figure 3.**
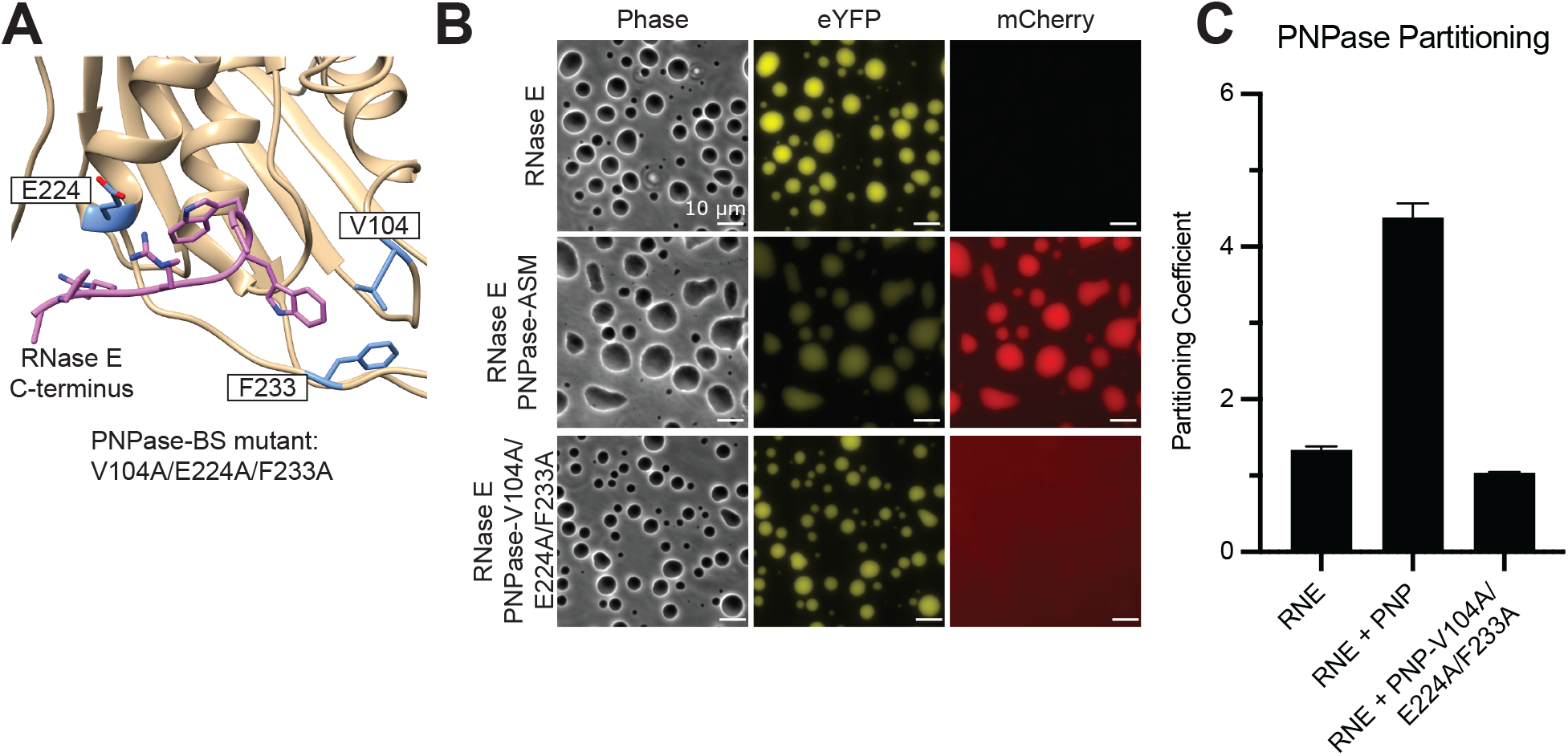
Identification of PNPase residues critical for interaction with RNase E. (A) Co-crystal structure of *C. crescentus* RNase E (PDB 4AIM)^30^ bound to PNPase highlights the critical protein-protein interaction site. (B) Phase contrast and fluorescence microscopy images of RNase E biomolecular condensates mixed with PNPase-ASM-mCherry or the PNPase-V104A/E224A/F233A-mCherry variant. The scale bar is 10 µm. (C) Average partitioning ratios with standard deviations are presented for PNPase and the PNPase-V104A/E224A/F233A-mCherry variant. Amongst experiments, PNPase is only significantly recruited into RNase E biomolecular condensates (p < 0.001). The PNPase-V104A/E224A/F233A-mCherry variant does not significantly partition into RNase E biomolecular condensates (p > 0.05). Data represent the average and standard deviation of three replicates.

We next considered if the short peptide representing the 10 C-terminal residues of RNase E could disrupt PNPase association with RNase E (EKPRR<u>GWW</u>RR)^24^. From here on, we refer to this as the GWW peptide. We found that the GWW peptide did not disrupt RNase E phase separation (Figure S2A, S2B). In contrast, PNPase partitioning decreased from 2.8 ± 0.2 to 1.7 ± 0.1 with the addition of 100 µM GWW peptide (Figure S2C). Interestingly, the high levels of GWW peptide resulted in a patchy appearance of PNPase-mCherry within the RNase E protein-rich droplets. The peptide outcompetes the C-terminus of RNase E for binding with PNPase, lowering the amount of PNPase associated with the BR-bodies. These results suggest that short peptides that can compete with RNase E for interaction with PNPase could function as inhibitors of PNPase recruitment into RNase E biomolecular condensates.

### RNA is not sufficient for PNPase recruitment into RNase E biomolecular condensates

We next considered if RNA clients of RNase E can recruit additional RNA binding proteins that do not directly associate with RNase E. Such recruitment of non-clients would impact a BR-body’s capabilities to control their composition to facilitate mRNA decay versus other RNA modification biochemistry. Given that both RNase E and PNPase can bind RNA^30^, we interrogated whether poly(A), a preferred substrate for PNPase, could promote PNPase association with BR-bodies lacking the PNPase binding site. We found that poly(A) was insufficient to drive PNPase accumulation in RNase E biomolecular condensates lacking the C-terminal binding site at poly(A) concentrations ranging from 25 - 100 ng/µL (Figure S3 A-C).

These results suggest that poly(A) has a poor capacity to recruit in PNPase and that protein-protein interactions are likely the main driver of protein recruitment into BR-bodies.

### Magnesium and phosphate impact RNase E droplet formation in vitro

We next considered how the substrates, cofactors, and products of the PNPase might affect the phase properties of RNase E. For example, our past studies have shown that sodium chloride concentration above 250 mM prevents RNase E’s homotypic phase separation^20^. PNPase requires magnesium and phosphate to facilitate its exoribonuclease activity (Figure 4A), which may also impact the phase diagram of RNase E. Notably, magnesium phosphate displays poor solubility in water at about 2-50 mM, dependent upon salt and pH conditions. Therefore, we considered how magnesium and phosphate impact RNase E phase separation.

**Figure 4.**
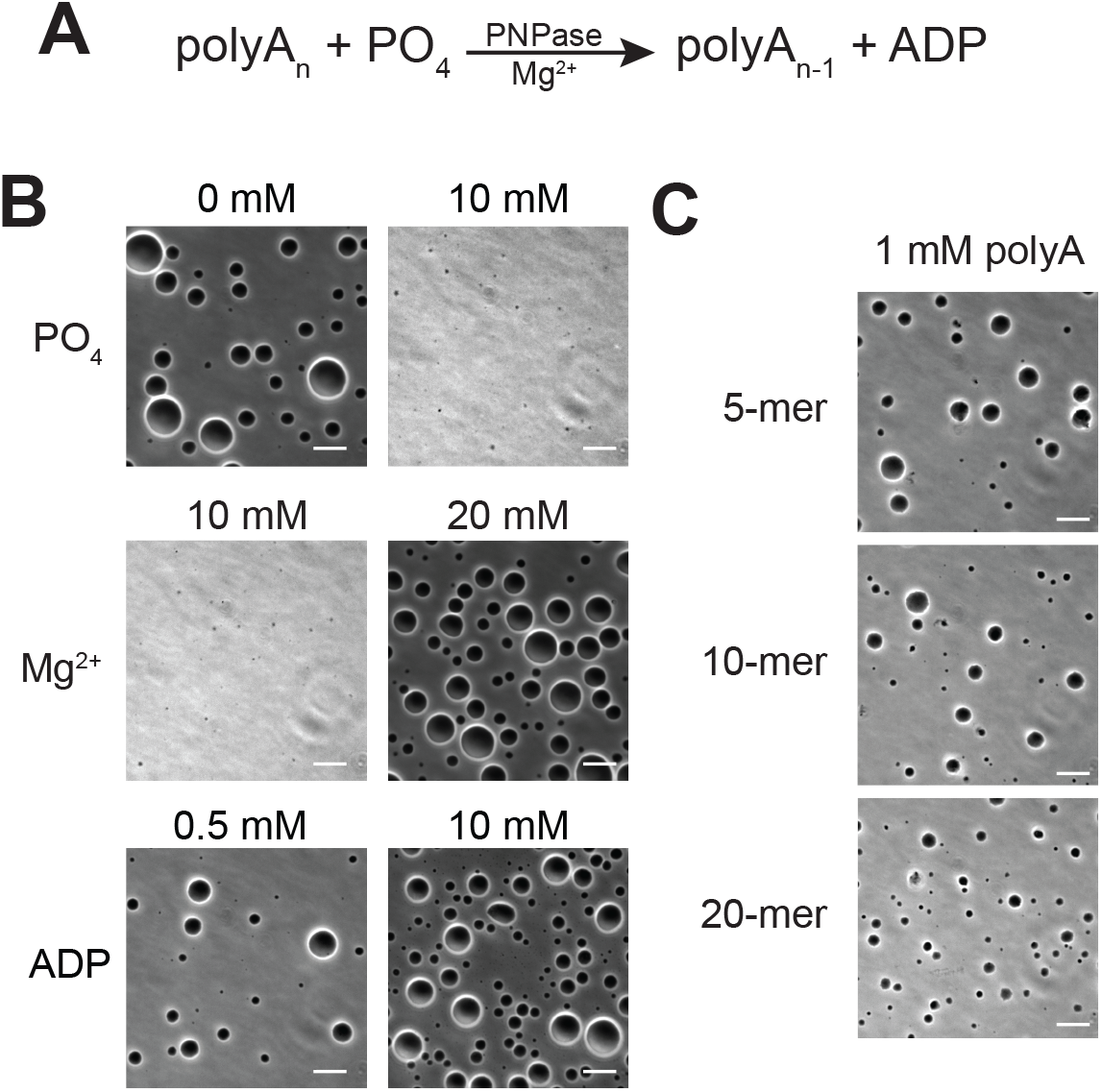
High sodium phosphate levels or low magnesium chloride levels dissolve the RNase E biomolecular condensates. Products of PNPase nuclease reaction do not dissolve biomolecular condensates. (A) PNPase and a magnesium ion cofactor catalyze the phosphorolysis of a single nucleotide (AMP) by adding inorganic phosphate to release nucleoside diphosphates (ADP). (B) Phase contrast images of RNase E biomolecular condensates in 0 mM or 10 mM sodium phosphate, 10 mM or 20 mM magnesium chloride, and 0.5 mM or 10 mM ADP. (C) Phase contrast images of RNase E biomolecular condensates mixed with 1 mM poly(A) 5-mer, 10-mer or 20-mer.

Above a critical concentration of 12 mM MgCl_2_, we observed 20 µM CTD-YFP phase-separated into biomolecular condensates (Figure 4B, Figure S4A). In contrast, we found that in the presence of 20 mM MgCl_2_, the addition of phosphate above 10 mM dissolved the RNase E biomolecular condensates (Figure 4B, S4A). Thus, the RNase-PNPase complex at 20 µM CTD-YFP forms a protein-rich phase in 20 mM MgCl_2_ with sodium phosphate levels that range from 0-8 mM (Figure S4A). This provides reaction conditions to characterize PNPase enzymatic function in which CTD-YFP forms robust protein-rich biomolecular condensates. In addition, the sensitivity of RNase E phase separation to magnesium and phosphate suggests that magnesium and phosphate cytosolic concentrations may impact BR-body formation *in vivo*.

### BR-body degradation products do not dissolve RNase E biomolecular condensates

A second consideration is if the products of PNPase exoribonuclease activity impact RNase E’s phase separation capabilities. Given that PNPase’s ribonuclease activity results in the production of NDPs, we examined if ADP could dissolve the RNase E biomolecular condensates *in vitro*. We found that the addition of ADP from 0.5 to 10 mM did not dissolve the RNase E-eYFP biomolecular condensates (Figure 4B, S4B). This indicates that ADP products of PNPase nuclease activity do not directly regulate RNase E’s phase separation. The ability to avoid dissolution at high levels of ADP is in contrast to observations by Saurabh *et al*. that showed that ATP could dissolve SpmX biomolecular condensates at concentrations at and above 2 mM^31^. This suggests that biomolecular condensation phase diagrams display varying robustness to ATP. Interestingly, in contrast to SpmX, RNase E may be responsive to phosphate nutrients directly instead of ATP or ADP (Figure 4B and S4).

We next considered if short oligoribonucleotides may be more effective at dissolving RNase E droplets than ADP (Figure 4C, S4C). It is possible that PNPase stalled on a reaction substrate could release a short oligonucleotide product. Poly(A) oligos of lengths 5, 10, and 20 nucleotides were incubated at 500 µM and 1000 µM with 20 µM RNase E to examine how short RNA products affect RNase E phase separation properties (Figure 4C, S4C). None of these poly(A) oligos at these concentrations caused the dissolution of RNase E droplets. These results indicate that the ribonuclease activity products of PNPase do not cause RNase E biomolecular condensates to dissolve.

### BR-bodies enhance PNPase nuclease activity against poly(A)

Past studies have shown that folded RNA substrates of PNPase require polyadenylation and the RNA helicase RhlB to promote robust PNPase exoribonuclease activity^26^. To decouple the effects of RhlB upon PNPase, we utilized poly(A) and poly(U) RNA as a PNPase substrate since they lack secondary structure. We mixed 20 µM CTD-YFP and 5 µM PNPase-mCherry with 25 ng/µL of poly(A) to assay PNPase’s exoribonuclease activity. After initiating reactions with poly(A), we tracked poly(A) RNA degradation using Urea PAGE gels stained with SYBR Gold RNA stain (ThermoFisher). In the absence of RNase E, PNPase degraded poly(A) at 80 ± 13 µg·min^-1^·(mg PNPase)^-1^ (Figure 5A). This rate of degradation by *C. crescentus* PNPase is about 10-fold lower than past reports of *E. coli* PNPase enzymatic functions^32^. These differences may be due to intrinsic activity differences between the two PNPase homologs or differences in the temperature during the assays (room temperature versus 37°C).

**Figure 5.**
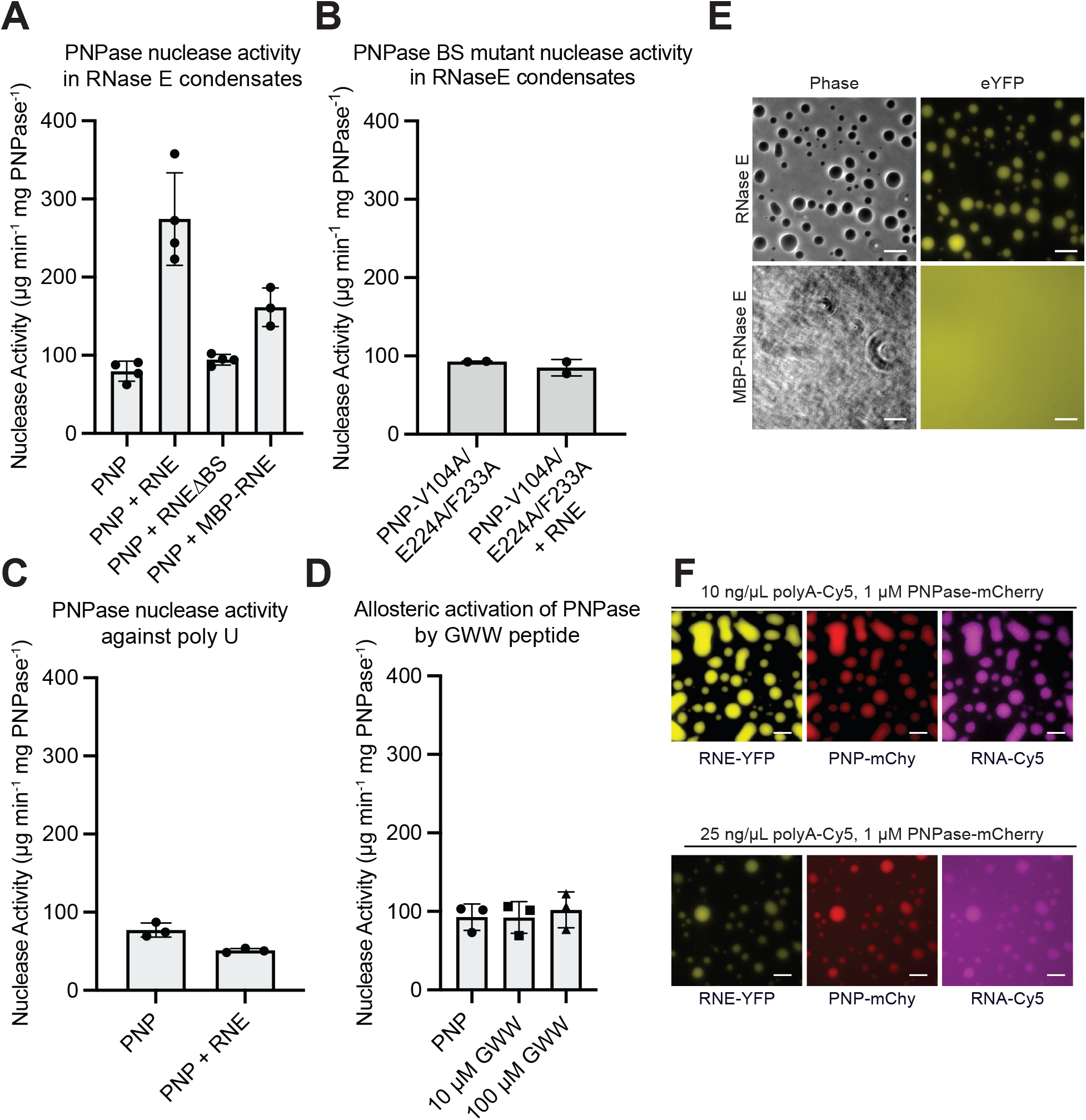
RNase E biomolecular condensates regulate the ribonuclease functions of PNPase. (A) Rate of ribonuclease activity towards 25 ng/µL poly(A) for 5 µM PNPase alone, 5 µM PNPase and 20 µM RNase E, and 5 µM PNPase and 20 µM RNase E-ΔBS (RNase E lacking the 14 C-terminal amino acids to which PNPase binds). (B) Ribonuclease activity of PNPase-V104A/E224A/F233A-mCherry (PNPase triple mutant lacking the ability to bind RNase E) and PNPase-V104A/E224A/F233A-mCherry mixed with RNase E. No significant rate increase was observed when RNase E was added (p > 0.05). Data are the average and standard deviation of three replicates. (C) Incorporation with BR-bodies does not stimulate PNPase’s decay of poly(U). Rate of ribonuclease activity towards 25 ng/µL poly(U) for 5 µM PNPase alone, or 5 µM PNPase and 20 µM RNase E. (D) The addition of 10 µM or 100 µM GWW peptide does not significantly alter the ribonuclease activity of 5 µM PNPase compared to PNPase alone (p > 0.05). (E) Fusion of maltose binding protein (MBP) significantly reduced the phase separation properties of RNase E. Phase contrast and fluorescence microscopy images of RNase E CTD-YFP versus RNase E MBP-CTD-YFP. (F) Phase contrast and fluorescence microscopy images of RNase E CTD-YFP mixed with PNPase-mCherry and poly(A)-Cy5 (upper) or poly(U)-Cy5 (lower). Droplets shown with poly(A)-Cy5 are induced by 10% PEG and appear weakly enriched with poly(A). Droplets shown with poly(U) are magnesium induced (0% PEG), where poly(U)-Cy5 recruits poorly into RNase E biomolecular condensates.

In contrast, the mixture of PNPase containing RNase E biomolecular condensates degraded poly(A) at a rate of 270 ± 60 µg·min^-1^·(mg PNPase)^-1^. This corresponds to a 3.4-fold enhancement of the poly(A) degradation rate over PNPase alone (p < 0.001) (Figure 5A). In comparison, RNase E-ΔPNP-BS had minimal effect on PNPase-mediated poly(A) degradation with a rate of 94 ± 7 µg·min^-1^·(mg PNPase)^-1^, a 1.2 fold enhancement which was not significantly different from WT-PNPase (p > 0.05) (Figure 5A). This indicates that non-interacting biomolecular condensates have little impact on PNPase functions.

While the addition of RNase E led to a 3.4-fold rate enhancement for wild-type PNPase, we found the addition of RNase E to the PNPase-V104A/E224A/F233A variant did not stimulate poly-A degradation. PNPase-V104A/E224A/F233A alone degraded poly(A) at a rate of 93 ± 1 µg·min^-1^·(mg PNPase)^-1^. In comparison, the PNPase-V104A/E224A/F233A variant mixed with RNase E degraded poly(A) at a rate of 85 ± 11 µg·min^-1^·(mg PNPase)^-1,^ which was not a significant difference in rate (p > 0.05) (Figure 5B). This bolsters the conclusion that a specific protein-protein interaction of RNase E’s C-terminal residues enhances PNPase’s ribonuclease activity.

### BR-bodies selectively enhance the degradation of poly(A) but not poly(U)

While PNPase can degrade any sequence of unstructured RNA, *in vivo* mRNA decay is stimulated by the presence of a 3’ polyadenylated tail^33^. Therefore, we investigated if the RNase E biomolecular condensates lead to an equal enhancement to the decay of other RNAs. Free PNPase degrades poly(U) at a rate of 77 ± 9 µg·min^-1^·(mg PNPase)^-1^, which is not significantly different from the rate at which PNPase degrades poly(A) (p > 0.05) (Figure 5C). Interestingly, we observed that the addition of RNase E to PNPase led to a decreased poly(U) degradation rate of 51 ± 3 µg min^-1^ mg PNPase^-1^ (Figure 5C). Therefore, unlike the 3-6 fold enhancement of poly(A) degradation mediated by RnaseE, we observed a 34% reduction in poly(U) degradation upon RNase E addition. These results indicate *in vitro* that RNase E may impart changes in substrate selectivity for PNPase.

In bacteria, polyadenylation of RNAs mediates their rapid decay. However, our past *in vivo* studies in *C. crescentus* indicated that the percent of A ribonucleotides is negatively correlated (R = -0.52) with BR-body enrichment^34^. This may suggest that long contiguous stretches of adenosine display preferences over isolated adenosine bases amongst oligoribonucleotide substrates. In addition, our past *in vivo* studies indicate that BR-bodies are preferentially associated with long and unstructured RNA substrates^34^. Like poly(A), poly(U) is also unstructured. Microscopy of RNase E droplets with Cy5-labeled poly(U) added confirms that poly(U) is only mildly enriched inside of RNase E droplets, with poly(U) partitioning ratios of 1.1 ± 0.1 without PNPase and 1.3 ± 0.1 with PNPase (Figure 5E). Therefore, the degree of RNA enrichment into the RNase E biomolecular condensates appears to regulate the degree of PNPase activity enhancement.

### RNase E’s scaffolding and phase separation play a key role in PNPase regulation

We considered three ways RNase E could stimulate PNPase activity towards poly(A). The first is that RNase E’s C-terminal binding site allosterically activates PNPase. In the second model, RNase E brings PNPase and poly(A) nearby via scaffolding, stimulating enhanced exoribonuclease activity. Finally, a third model considers the unique chemical environment of biomolecular condensates that concentrates both poly(A) and PNPase. This third model builds upon the scaffolding effect to include the impact of a higher concentration of PNPase and poly(A) in the BR-body, thereby increasing the kinetics of PNPase through mass action.

To test these models, we incubated PNPase with the GWW peptide from RNase E for 30 min before measuring exonuclease activity against poly(A). There was no statistically significant difference in PNPase activity when 10 or 100 µM of the peptide was added (p > 0.05) (Figure 5D), indicating that the RNase E GWW peptide does not allosterically activate PNPase.

To test the second model, we used a maltose-binding protein fusion of RNase E’s CTD, called MBP-RNase E. MBP-RNase E retains the RNA and PNPase binding sites but cannot phase separate *in vitro* (Figure 5F). The PNPase activity in the presence of MBP-RNase E was elevated 2-fold over PNPase alone to 160 ± 20 µg·min^-1^·(mg PNPase)^-1^ (p < 0.05) (Figure 5A). Analysis of these constructs suggests that scaffolding in the dilute phase increases PNPase function 2-fold. In comparison, phase separation of the RNase E-PNPase scaffolded complex yields a 3.4-fold enhancement and an additional 1.7-fold increase in PNPase activity over scaffolded PNPase (Figure 5A). Therefore, there are additive rate enhancements from scaffolding alone and mass action effects of phase separation.

## Discussion

This study investigated how RNase E regulates PNPase activity and how PNPase functions impact RNase E phase separation. We found that critical cofactors and substrates of PNPase, such as divalent magnesium and phosphate ions, regulated the phase separation properties of RNase E protein-rich biomolecular condensates *in vitro*. Interestingly, model system studies of arginine-rich peptides mixed with RNA showed that various divalent ions could alter the material properties and the switch from heterotypic to homotypic phase separation^35^. These results suggest that magnesium and phosphate nutrients may alter the material properties and composition of BR-bodies. In contrast, we found that the products of PNPase exoribonuclease activity had little impact on RNase E’s phase separation properties (Figure 4).

From past studies, it is well known that RNase E’s CTD domain regulates the functions of PNPase^24, 36^. Here we considered the mechanism of how RNase E regulates the exoribonuclease functions of PNPase. As a part of this work, we identified a PNPase variant that disrupts the RNase E-PNPase interaction, indicating a specific RNase E-PNPase interaction site mediates their recruitment into the RNase E biomolecular condensates. Moreover, this interaction leads to a 3.4-fold enhancement of PNPase’s enzymatic functions (Figure 5).

Our results suggest dual contributions from scaffolding RNA and PNPase near each other and from mass action effects of crowding several complexes in close proximity within a phase-separated environment. Specifically, the PNPase partitioning ratio was enriched nearly 6-fold over the dilute phase. Given that PNPase forms a trimer that degrades a single RNA substrate, one may expect to observe a 2-fold increase in poly(A) degradation due to mass action effects. Consistent with mass action effects, we observed a significant 1.7-fold enhancement (p < 0.01) in PNPase function inside BR-bodies over the function of PNPase in the non-phase separating MBP-CTD: PNPase complex (Figure 5A).

One limitation of biomolecular condensates is that the viscosity of the environment may limit the benefit of increased concentration. For example, one synthetic system observed the enzymatic activity of adenylate kinase was dampened within Dbp1N-AK-Dbp1C biomolecular condensates when compared to the increase in concentration^37^. In contrast, a synthetic system where the SUMOylation enzyme pathway was recruited into engineered condensates leveraged the full effects of mass action^38^. This suggests that the strength of interactions between the substrates, products, and scaffolds can lead to substrate exclusion or substrate enrichment and alter the viscosity of the biomolecular condensates, fine-tuning the mass transfer effects on enzyme performance.

For example, studies from Peeples *et al*. systematically considered how the SUMOlyation enzymatic cascade was regulated by biomolecular condensates^38^. A key observation was that changes in enzyme activity were due to scaffold-induced changes in substrate *K*_m_^38^. We compared poly(A) and poly(U) to explore how the RNase E-PNPase biomolecular condensates degrade RNAs of a varied sequence. Both poly(A) and poly(U) serve as suitable substrates for the free PNPase (Figure 5A, 5C). While the RNase E-PNPase biomolecular condensates stimulated a 3.4-fold stimulation of poly(A) degradation over PNPase alone, we observed that RNase E-PNPase biomolecular condensates led to mild repression of poly(U) degradation (Figure 5C). This decreased performance towards the poly(U) substrate may be due to decreased affinity for the RNase E-PNPase biomolecular condensate.

Alternatively, the poly(U) may alter the viscosity of the RNase E-PNPase biomolecular condensate and alter the potential benefits of mass action. This preference for poly(A) over poly(U) is intriguing, as polyadenylation has been implicated in mRNA degradation in *E. coli*^*39*^ and *C. crescentus*^*40*^. Indeed, results suggest that BR-bodies can fine-tune the half-lives of RNAs in cells and that polyadenylation may shape the available transcriptome. Therefore, future studies will be needed to examine the interplay of polyadenylation and BR-body functions *in vivo* and *in vitro*.

Beyond mass action impacts upon client enzymes, the interaction between scaffold and client may allosterically regulate enzyme functions. Here we showed that the addition of the GWW peptide did not stimulate changes in PNPase function (Figure 5D). This suggested that allosteric regulation likely does not play a major role in how BR-bodies impact PNPase’s enzymatic functions. In contrast, our past studies showed that the PodJ scaffold allosterically regulates the histidine kinase PleC through interaction with PleC’s sensory domain^17^. The coupling of enzyme recruitment and enzyme regulation enabled the spatial regulation of PleC function, which is critical for cell polarity. Overall, our studies suggest that the systematic evaluation of biomolecular condensate effects through selective scaffolding, mass action, and allostery can reveal how biomolecular condensates fine-tune enzyme functions.

## Methods

### Protein expression and purification

Purification of PNPase-mCherry: Plasmid pMJC0095 was constructed to express PNPase from *C. crescentus* with an N-terminal 6x-His tag and a C-terminal mCherry. Plasmid pTEV5-PNPase-mCherry was transformed into chemically competent Rosetta (DE3) cells and plated onto LB-Miller plates supplemented with 50 mg/mL chloramphenicol, 100 mg/mL ampicillin and incubated overnight at 37 °C. From a single colony, an overnight 60 mL LB-Miller culture (30 mg/mL chloramphenicol, 50 mg/mL ampicillin) was inoculated and incubated at 37°C. From this saturated culture, 6 L of LB-Miller media (30 mg/mL chloramphenicol, 50 mg/mL ampicillin) was inoculated with 6 mL of the saturated culture and grown to mid-log phase (∼0.5 OD600). Expression of PNPase-mCherry was induced with 333 µM isopropyl-β-D-1-thiogalactopyranoside (IPTG) for 4 h at 25 °C. The cells were collected by centrifugation at 4 °C, 4000 g, for 30 min. The resulting pellet was washed with 60 mL resuspension buffer (50 mM Tris pH 7.5, 500 mM NaCl) before being pelleted again at 4 °C, 4000 g, for 20 min and stored at -80 °C.The cell pellet was thawed on ice and then resuspended in 10 mL lysis buffer per liter of culture (20 mM Tris HCl pH 7.5, 500 mM NaCl, 5 mM imidazole, 200 U benzonase) supplemented with SigmaFast protease inhibitor tablets (Sigma). The cell suspension was lysed by continuous passage through an Avestin Emulsiflex-C3 at 15,000 psi for 15 min at 4 °C. Cell debris was pelleted by centrifugation at 29,000 g for 45 min at 4 °C. The supernatant was loaded onto a HisTrap FF column (GE Healthcare) and washed with 20 column volumes of wash buffer (20 mM Tris HCl pH 7.5, 500 mM NaCl, 5 mM imidazole). Then was eluted with elution buffer (20 mM Tris HCl pH 7.5, 500 mM NaCl, 500 mM imidazole). Fractions containing PNPase were supplemented with 50 mM sodium phosphate (pH 7.5) at 37 °C for 1 h to drive phosphorolysis of co-purifying RNA. The fractions were loaded onto a G-Sep™ 6-600 kDa Size Exclusion Columns (G-Biosciences) and eluted with storage buffer (20 mM Tris HCl pH 7.5, 500 mM NaCl, 5% (v/v) glycerol). Fractions containing PNPase were concentrated using 50,000 MWCO Amicon centrifugal filters to 15.5 mg/mL, aliquoted, and flash frozen in liquid nitrogen and stored at -80 °C.

Purification of RNase E-CTD-YFP: RNase E CTD was purified as described previously^41^, and summarized here. Cells were thawed on ice and then resuspended in 10 mL lysis buffer per liter of culture (20 mM Tris HCl pH 7.5, 1 mM NaCl, 20 mM imidazole, 1mM beta-mercaptoethanol, 20 U DNase I, 100 U RNaseA, 200 U benzonase, 0.1% Triton X-100) supplemented with SigmaFast protease inhibitor tablets (Sigma). The cell suspension was lysed by continuous passage through an Avestin Emulsiflex-C3 at 15,000 psi for 15 min at 4 °C. Cell debris was pelleted by centrifugation at 29,000 g for 45 min at 4 °C. The supernatant was loaded onto a HisTrap FF column (GE Healthcare) and washed with 20 column volumes of wash buffer (20 mM Tris HCl pH 7.5, 1000 mM NaCl, 20 mM imidazole, 1 mM beta-mercaptoethanol). Subsequently, the purification was eluted with elution buffer (20 mM Tris HCl pH 7.5, 1000 mM NaCl, 500 mM imidazole, and 1 mM beta-mercaptoethanol). Fractions containing RNase E were loaded onto a G-Sep™ 6-600 kDa Size Exclusion Columns (G-Biosciences) and eluted with storage buffer (20 mM Tris HCl pH 7.5, 500 mM NaCl, 1 mM beta-mercaptoethanol). Fractions containing RNase E were concentrated using 50,000 MWCO Amicon centrifugal filters to 18.6 mg/mL, aliquoted, flash frozen in liquid nitrogen and stored at -80 °C.

MBP-RNase E-CTD-eYFP was purified the same way as RNase E-CTD-eYFP except for an additional nucleic acid removal step. After MBP-RNase E-CTD-eYFP was eluted from the HisTrap column, the protein was desalted using a PD-10 column (GE healthcare) with heparin column binding buffer (20 mM Tris HCl pH 7.5, 1 mM beta-mercaptoethanol, 10% glycerol) before passage over a heparin column (Cytiva). MBP-RNase E-CTD-eYFP was eluted with Heparin elution buffer (20 mM Tris HCl pH 7.5, 1 mM beta-mercaptoethanol, 10% glycerol, 2 M NaCl) using a linear gradient. Protein was then buffer exchanged into storage buffer (20 mM Tris HCl pH 7.5, 200 mM NaCl, 1 mM beta-mercaptoethanol, 10% glycerol) using a PD-10 column, concentrated using 50,000 MWCO Amicon centrifugal filters to 23.5 mg/mL, aliquoted, and flash frozen in liquid nitrogen before storage at -80 °C.

### Microscopy

Fluorescence microscopy samples were prepared by thawing requisite proteins on ice and mixing a buffer to create a final concentration of 20 mM Tris pH 7.5, 1 mM MgCl^2^, 10 mM Na^2^HPO^4^ pH 7.5, 100 mM NaCl, 10% PEG 8000 (Figures 2, S1, S2 [120 mM NaCl], S3) or 20 mM Tris pH 7.5, 20 mM MgCl^2^, 4 mM Na^2^HPO^4^ pH 7.5, 70 mM NaCl, 0.5 mM DTT (Figures 3, 4, 5, S4, S5), to which was added protein and poly(A) at the necessary concentrations specified in each experiment. Imaging samples were pipetted into a 1 mm adhesive spacer (Electron Microscopy Sciences) affixed to a microscope slide (VWR) and sealed with a glass coverslip (VWR). Slides were inverted and allowed to sit at room temperature for 30 minutes before imaging on a Nikon Eclipse Ti-E inverted microscope with a Plan Apo-(lambda) 100x/1.45 oil objective and 518F immersion oil (Zeiss). Excitation filter cubes CFP/YFP/mChy (77074157) and Cy5 (77074160) from Chroma and emission filter sets CFP/YFP/mChy (77074158), and Cy5 (77074161) from Chroma were used for fluorescence imaging with a Spectra X light engine from Lumencor. Images were taken with an Andor Ixon Ultra 897 EMCCD camera.

Images were analyzed with Fiji using a gaussian blur image subtraction and Renyi Entropy threshold method to find droplet boundaries. The mean signal from each droplet area was divided by the mean signal of the non-droplet area to give a partitioning ratio for each droplet, which is averaged to give a partitioning ratio for each experimental condition.

### Generation of fluorescent nucleotides

Fluorescent polynucleotides were generated using 5 µM PNPase with 99 µM NDP and 1 µM ADP-Cy5 in a buffer of 20 mM Tris pH 7.5, 100 mM NaCl, 10 mM MgCl^2^, 0.5 mM DTT. Reactions were run for 2 h at room temperature. Fluorescent polynucleotides were purified using silica gel spin columns and frozen at -80 °C for future use.

### PNPase mediated RNA-degradation assay

RNA degradation assays were performed at room temperature in 20 mM Tris-HCl (pH 7.5), 70 mM NaCl, 20 mM MgCl^2^, 4 mM Na^2^HPO^4^ (pH 7.5), and 0.5 mM DTT with 5 µM purified PNPase and 20 µM purified RNase E. Reactions were initiated by adding 25 ng/µL poly(A) RNA. For time course assays, aliquots were withdrawn and quenched in 100 mM EDTA. Samples were denatured in 1.5 volumes of 2x RNA loading buffer containing 95% formamide, 18 mM EDTA, and 0.025% SDS and incubated at 95 °C for 3 min. Quenched ribonuclease reaction aliquots were loaded onto a pre-run 6% acrylamide gel containing 7 M urea. The gel was run in 1x TBE (89 mM Tris base, 2 mM EDTA, 89 mM boric acid) at 250 V at room temperature to separate RNA. Subsequently, the gel was rinsed in Milli-Q water for 5 minutes and stained for 20 minutes with 1x SYBR gold nucleic acid stain (Invitrogen) in 1x TBE. Each gel assay included RNA only and protein only controls. Gels were imaged with BioRad ChemiDoc™ MP imager using SYBR gold settings and quantified using the BioRad ImageLab software package. The intensity of the protein-only lane was subtracted from each timepoint lane intensity and plotted against time. The degradation rate was calculated in relation to a known amount of RNA added in an RNA-only control lane and divided by the amount of protein to give rates in µg·min^-1^·(mg PNPase)^-1^. Since poly(A) is of heterogenous length, nuclease activity cannot be reported on a molar scale.

This is a test

### PNPase mediated RNA-degradation assay with GWW peptide

The 10 C-terminal residues of *Caulobacter crescentus* RNase E (EKPRRGWWRR) (GWW peptide) were synthesized by GenScript with C-terminal amidation and dissolved in Milli-Q water, and frozen at -80 °C until further use. In RNA-degradation assays containing GWW peptide, the peptide was added to the reaction tube with buffer and PNPase and incubated at room temperature for 30 minutes to allow the peptide to associate with PNPase. Reactions were otherwise carried out as described previously.

## Notes

### Competing Interest Statement

The authors have declared no competing interest.

