## Supporting Information for "RNase E biomolecular condensates stimulate PNPase activity"

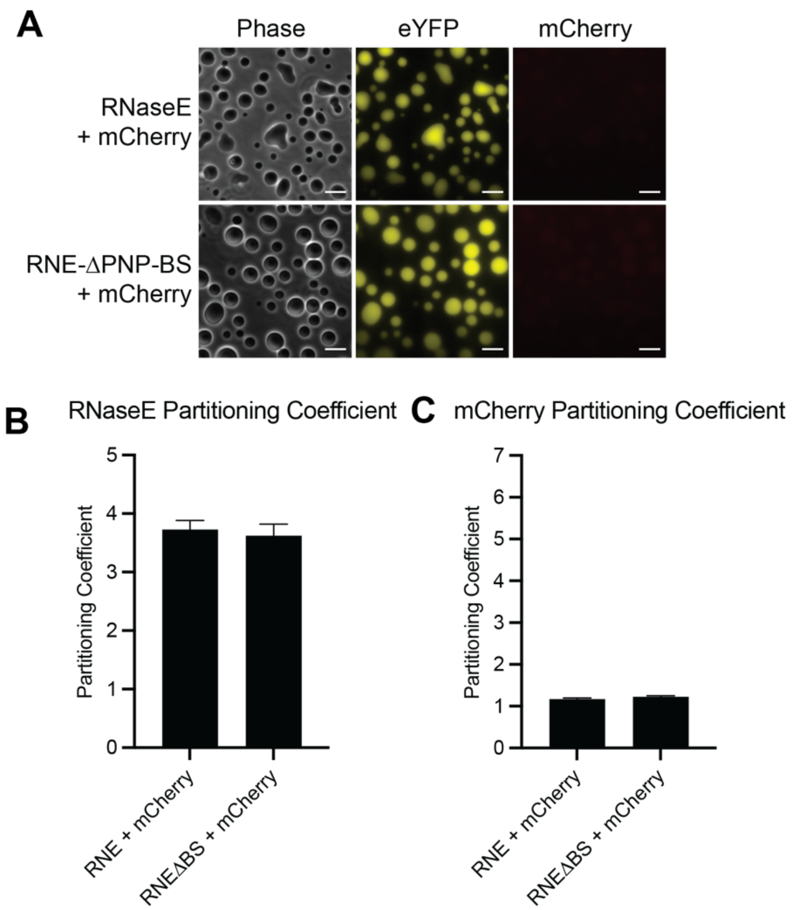


**Figure S1: Weak fluorescent protein interactions between eYFP and mCherry do not lead to recruitment of mCherry into RNase E-eYFP or RNE-ΔPNP-BS protein-rich biomolecular condensates**. (A) Phase contrast and fluorescence microscopy images of 20 μM RNase E

biomolecular condensates mixed with 1 μM mCherry. RNaseE and RNaseE-ΔPNP-BS contain a C-terminal eYFP tag. The scale bar is 10 μm. (B) Average partitioning ratios with their standard deviations are presented for RnaseE-eYFP and RNaseE-ΔPNP-BS-eYFP. There is no significant difference between any PRs ($p > 0.05$). (C) Average partitioning ratios with their standard deviations are presented for mCherry. There is no significant difference between any PRs ($p > 0.05$). Data represent average and standard deviations of $n > 300$ droplets.

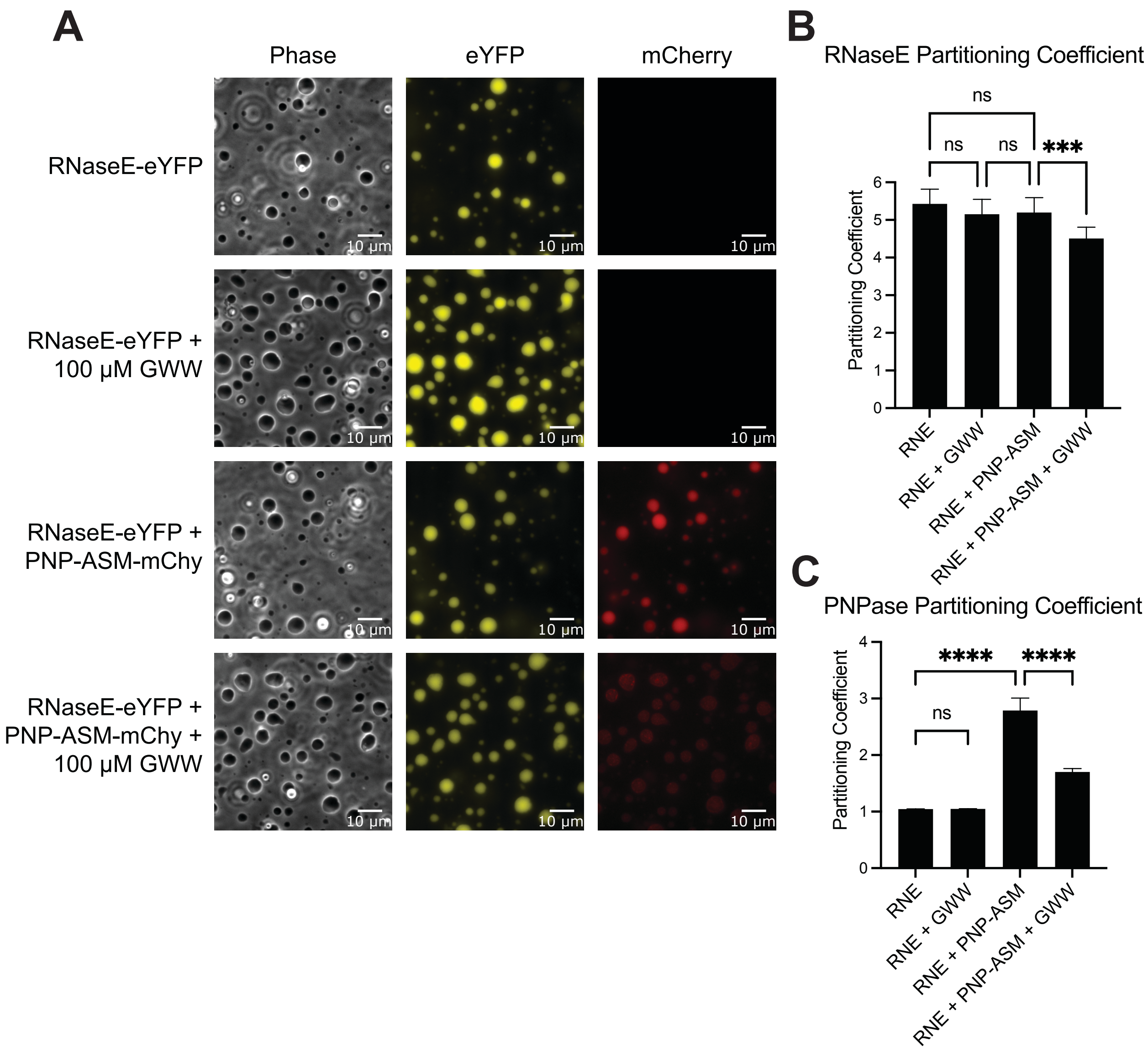


**Figure S2: The RNase E GWW peptide can chase PNPase-ASM-mCherry out of RNase E biomolecular condensates** (A) Phase contrast and fluorescence microscopy images of protein-rich biomolecular condensates formed by RNase E-eYFP were incubated with a peptide comprised of the 14 C-terminal residues of RNase E (called the GWW peptide). A mixture of 1 µM PNPase together with 100 µM of the GWW peptide was incubated for 60 minutes. After this incubation, the mixture was added to 20 µM of RNase E biomolecular condensates. Buffer conditions were 20 mM Tris pH 7.5, 120 mM NaCl, 1 mM $MgCl_2$, 10 mM $NaPO_4$ pH 7.5, 10% PEG. Partitioning coefficients of RNase E-eYFP (B) and PNPase-ASM-mCherry (C) from three replicates containing over 300 droplets. The lower partitioning ratio for PNPase is likely due to the 120 mM NaCl concentration which is higher than was used in other imaging experiments. GWW peptide does not significantly alter RNase E droplet formation or partitioning ratios. However, it does significantly decrease the partitioning of PNPase into the droplets.

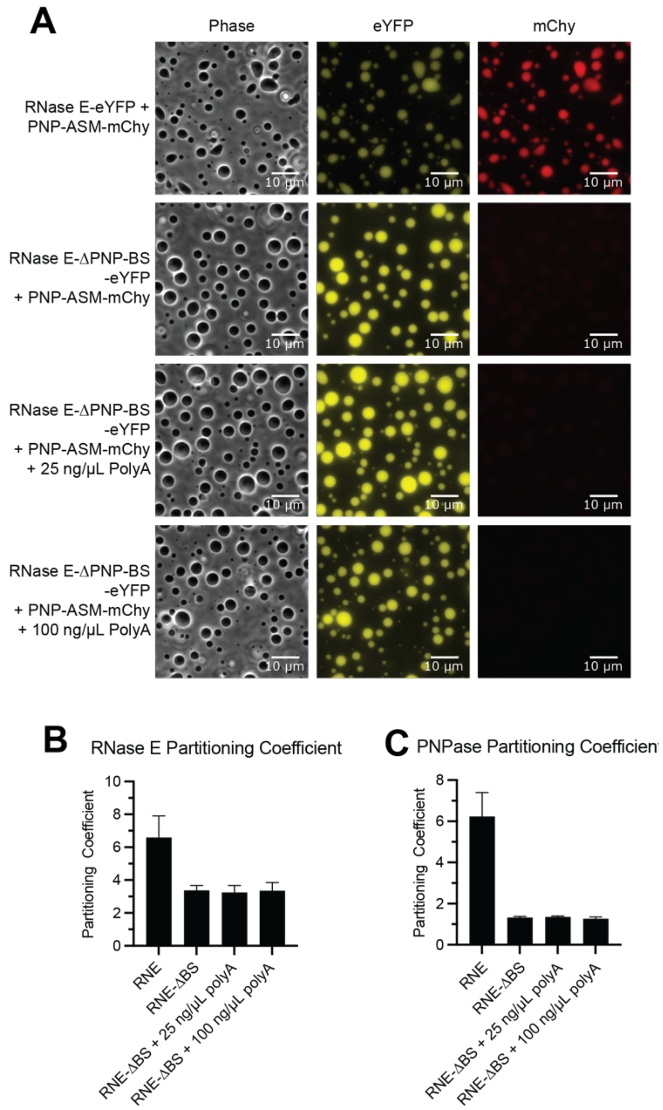


**Figure S3: Poly(A) RNA is insufficient to shepherd PNPase into BR-bodies formed by RNase E-ΔBS.** (A) Phase contrast and fluorescence microscopy images of RNase E-rich or RNase E-ΔBS-rich biomolecular condensates were incubated with 5 μM of PNPase-ASM-mCherry. 25 or 100 ng/μL poly(A) RNA was added to droplets formed with RNase E-ΔBS did not show incorporation of PNPase. Average and standard deviations of partitioning coefficients of RNase E (B) and PNPase (C) from over 300 droplets show a significant decrease in PNPase partitioning into condensates formed with RNase E-ΔBS even in the presence of poly(A) RNA.

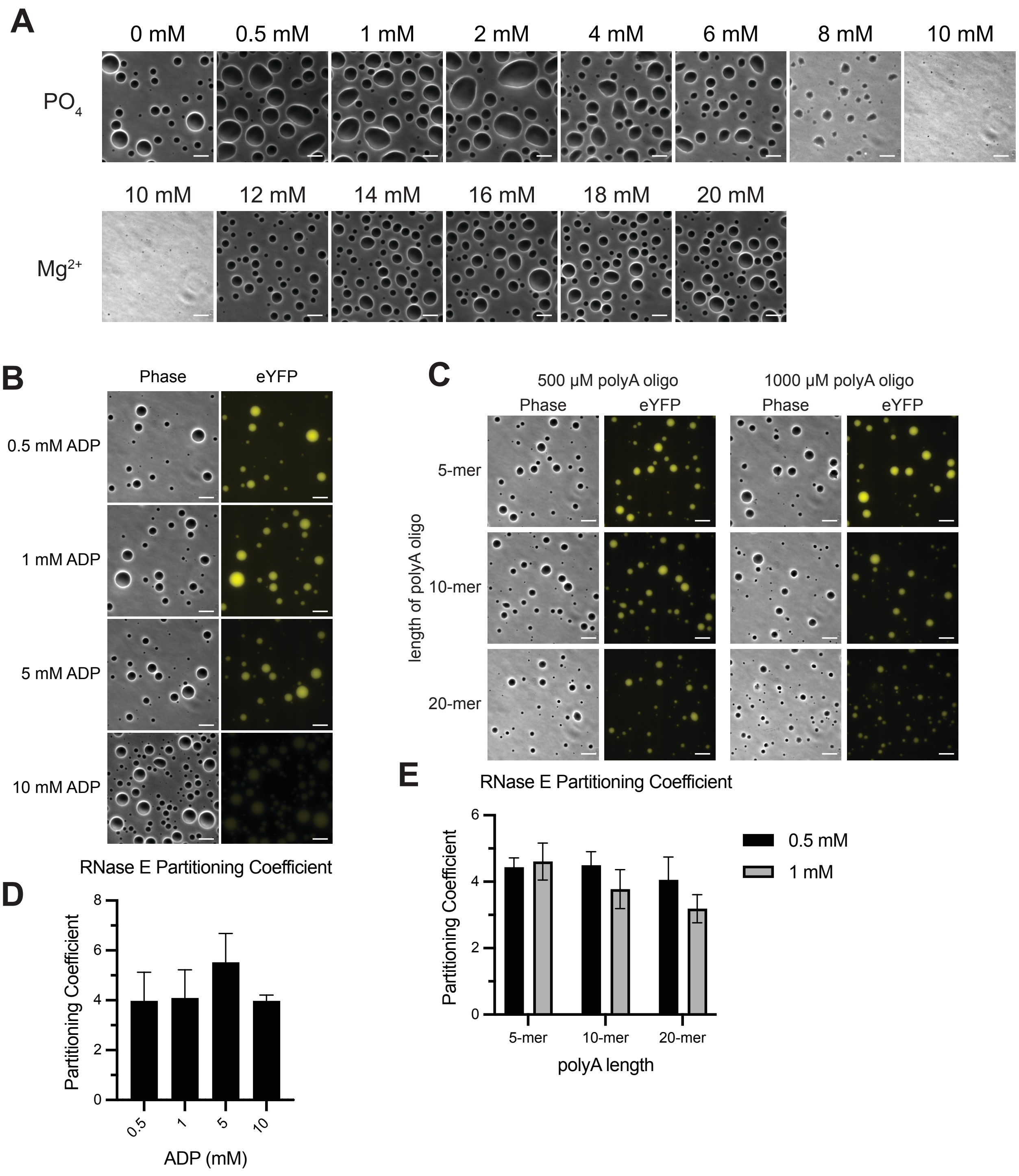


**Figure S4: Sodium phosphate and magnesium chloride regulate the formation of RNase E CTD biomolecular condensates.** (A) Phase contrast images of 20 µM of RNase E in 20 mM $MgCl_2$ mixed with 0-10 mM sodium phosphate. Phase contrast images of 20 µM of RNase E in 4

mM sodium phosphate mixed with 0-20 mM magnesium chloride. (B) ADP at concentrations of 0-10 mM was added to a solution of 20 μM of RNase E-eYFP and subsequently imaged by phase contrast and fluorescence microscopy. (C) Short poly(A) oligoribonucleotides (5-mer, 10-mer or 20-mer) were added to a solution of 20 μM of RNase E-eYFP and subsequently imaged by phase contrast and fluorescence microscopy. Scalebar = 10 μm. (D) Partitioning coefficients of RNase E-eYFP during ADP titration. (E) Partitioning coefficients of RNase E-eYFP with short poly(A) oligomers. Partitioning coefficients represent the average and standard deviation of n > 150 droplets.

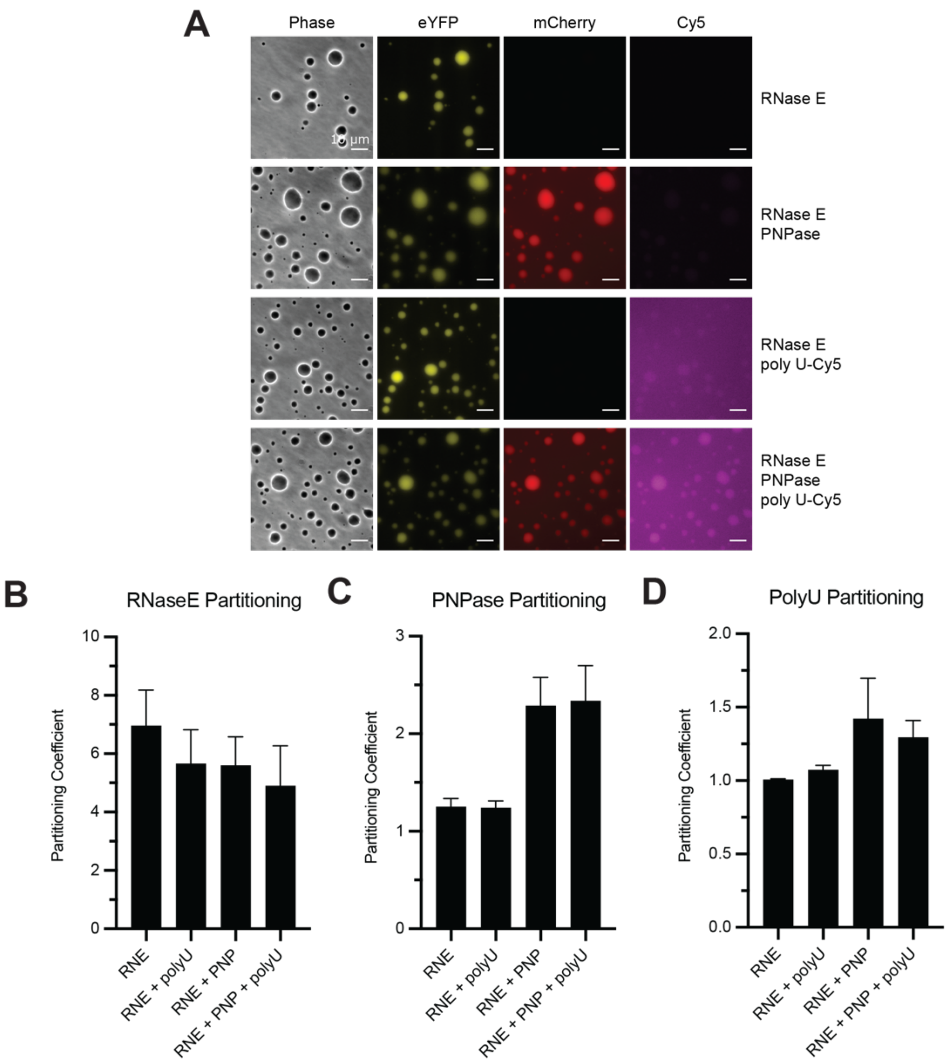


**Figure S5: Poly(U) is poorly enriched in RNase E biomolecular condensates.** (A) Phase contrast and fluorescence microscopy images of 20 μM of RNase E-eYFP, 20 μM of RNase E-eYFP with 5 μM PNPase, 20 μM of RNase E-eYFP with 25 ng/μL poly(U), and 20 μM of RNase E-eYFP with 5 μM PNPase and 25 ng/μL poly (U). Scalebar = 10 μm. Partitioning coefficients for (B) RNase E or (C) PNPase and (D) Poly (U) are calculated as the average and standard deviation of n > 300 droplets.

**Supplemental Materials, Methods, and procedures**

***Plasmids, primers, and strains***

Table 1. Plasmids used in this study

| Plasmid | Description | Reference |
|---|---|---|
| pTEV5 | Bacterial expression vector | [1] |
| pMJC0094 | His6x-PNPase | This study |
| pMJC0095 | His6x-PNPase-mCherry | This study |
| pMJC0112 | His6x-PNPase-S339A/S340A/S341A active site mutant | This study |
| pMJC0113 | His6x-PNPase-S339A/S340A/S341A-mCherry active site mutant | This study |
| pMJC0119 | His6x-RNase E(CTD)-Δ884-898-eYFP PNPase binding site deletion | This study |
| pMJC0137 | His6x-PNPase-V104A/E224A/F233A-mCherry | This study |
| pDT177 | His6x-pTEV5-RNase E(CTD)-eYFP | This study |
| pDT279 | His6x-pTEV5-MBP-RNase E(CTD)-eYFP | This study |

Table 2. Strains were used in this study.

| Strain | Description | Reference |
|---|---|---|
| E. coli DH5α | Bacterial cloning strain | Invitrogen |
| E. coli Rosetta (DE3) pLysS | Bacterial expression strain | Novagen |
| MJC192 | Rosetta-pMJC0094 PNPase expression strain | This study |
| MJC196 | Rosetta-pMJC0095 PNPase-mCherry expression strain | This study |
| MJC212 | Rosetta-pMJC0112 PNPase-S339-341A expression strain | This study |
| MJC213 | Rosetta-pMJC0113 PNPase-S339-341A expression strain | This study |
| MJC235 | Rosetta-pMJC0119 RNase E(CTD)-ΔPNPase_BS-eYFP expression strain | This study |
| MJC303 | Rosetta-pMJC0137 PNPase-V104A/E224A/F233A expression strain | This study |
| DTT210 | Rosetta-pDT279 MBP-RNase E(CTD)-eYFP expression strain | This study |

Table 3. Primers used in this study. The primer sequence is listed from 5' to 3'.

| Primer | 5'-3' sequence | Description |
|---|---|---|
| MJC222 | GAAAACCTGTATTTTCAGGGCGCTATGTTCGAT ATCAAACGCAAGACGATCGAGTGG | PNPase forward |
| MJC223 | GCTCGAGAATTCCATGGCCATATGGCTTTACGC CTCTTCGGCCGCCG | PNPase reverse |
| MJC224 | CTCGAGATCTTAACGCCTCTTCGGCCGCCGCTT CC | PNPase-mCherry reverse |
| MJC225 | GGCCGAAGAGGCGTTAAGATCTCGAGCTCCGG AGAATTCG | mCherry forward |

| | | |
|---|---|---|
| MJC226 | GCTCGAGAATTCCATGGCCATATGGCTTTACTT GTACAGCTCGTCCATGCCGCC | mCherry reverse |
| MJC261 | AGACCGTGGCCATCGCCGCCGCACCGTTGCTC TCGGTGATCTCCG | PNPase 1-338 (S339-441A) active site mutant reverse |
| MJC262 | CGGTGCGGCGGCGATGGCCACGGTCTGCGGT TCG | PNPase 442-713 active site mutant forward |
| MJC265 | CCAACTAGTGAAAACCTGTATTTTCAGGGCGCT ATGACCGGCGTGCTGGAAGG | RNase E(CTD) forward |
| MJC269 | GCTCGAGAATTCCATGGCCATATGGCTTTACTT GTACAGCTCGTCCATGCCCA | EYFP reverse |
| MJC280 | CGGAGCTCGAGATCTTAAGATCTCGTTCGGATC CGGCTCG | RNE-ΔPNP_BS reverse |
| MJC281 | GGATCCGAACGAGATCTTAAGATCTCGAGCTCC GGAGAATTCG | EYFP forward |
| MJC282 | TGTCTTCCGGCTCCGCCGCGAAGGGCTCCTTG GCGG | PNPase-F233A reverse |
| MJC283 | GCCCTTCGCGGCGGAGCCGGAAGACACCGAC GCG | PNPase-F233A forward |
| MJC290 | TCTTGAAGCCCTTCGCGAACAGCGGGCGGATC GGAC | PNPase-V104A reverse |
| MJC291 | CCCGCTGTTCGCGAAGGGCTTCAAGAACGAAG TC | PNPase-V104A forward |
| MJC292 | GCGGCGTGCGCGGCCAGGTCGATGATCGCGT CG | PNPase-E224A reverse |
| MJC293 | ATCATCGACCTGGCCGCGCACGCCGCCAAGGA GCCCTTCG | PNPase-E224A forward |
| DT471 | TTTTCAGGGCGCTAAAATCGAAGAAGGTAAACT GGTAATCTGG | MBP-forward |
| DT473 | GCACGCCGGTCATACCTTGGAAGTAGAGATTCT CTGACGTGG | MBP-reverse |
| DT476 | CTACTTCCAAGGTATGACCGGCGTGCTGGAAG G | RNase E(CTD)-eYFP forward |
| DT477 | TGGCCATATGGCTTTACTTGTACAGCTCGTCCA TGCCG | RNase E(CTD)-eYFP reverse |

***Plasmid construction procedures***

**Plasmid pMJC0094, PNPase expression vector:** pMJC0094 was generated in pTEV5 using the Gibson DNA assembly method [1, 2]. Primers for Gibson reactions were designed using J5 DNA assembly design automation software [3]. The resulting plasmid encodes a hexahistidine tag, followed by a TEV protease cleavage site at the N-terminus of PNPase. Full-length *C. crescentus* PNPase encoding residues 1-713 was amplified by PCR using genomic DNA of *Caulobacter crescentus* NA1000 as a template with primers MJC222 and MJC223. These primers include overhang sequences homologous to regions flanking the NheI site in pTEV5. Vector pTEV5 was linearized by digestion with NheI restriction enzyme and ligated with PCR fragments following the Gibson method described here briefly. Equimolar amounts of linearized pTEV5 and PNPase PCR fragments were incubated at 50 °C for 60 min in a reaction mixture containing 5% PEG-8000, 100 mM Tris-HCl pH 7.5, 10 mM $MgCl_2$, 10 mM DTT, 0.2 mM of each of the four dNTPs, 1.25 mM NAD, 5.3 mU/µL T5 exonuclease, 33.3 mU/µL Phusion DNA polymerase, and 5.3 U/µL Taq ligase (NEB) resulting in plasmid pMJC0094 that encodes an N-terminal $His_6$-tag and TEV cleavage site (MSYYHHHHHHDYDIPTSENLYFQGAM) fused to PNPase.

**Plasmid pMJC0095, PNPase-mCherry expression vector:** pMJC0095 encoding PNPase with a C-terminal flexible linker and mCherry was generated in pTEV5 using the Gibson assembly method. PCR fragments encoding full-length PNPase were cloned from pMJC0094 using primers MJC222 and MJC224. PCR fragment encoding a linker (LRSRAPENSNVTRHRSAT) and mCherry was cloned from pDT027 using primers MJC225 and MJC226. Primers MJC222 and MJC226 include overhang sequences homologous to regions flanking the NheI site in pTEV5. Gibson assembly was completed as previously described resulting in plasmid pMJC0095 that encodes an N-terminal $His_6$-tag and TEV cleavage site (MSYYHHHHHHDYDIPTSENLYFQGAM) fused to PNPase with a C-terminal flexible linker (LRSRAPENSNVTRHRSAT) and mCherry.

**Plasmid pMJC0112, PNPase-mCherry active site mutant expression vector:** pMJC0112 encoding PNPase-S339-341A with an active site mutant generated by serine to alanine triple mutation at residues 339-341 was generated in pTEV5 using the Gibson assembly method with primers designed by j5 DNA assembly design software. Codons 1-345 of PNPase were amplified by PCR from pMJC0094 with primers MJC222 and MJC261. Codons 337-713 of PNPase were amplified by PCR from pMJC0094 with primers MJC262 and MJC223. Codons 337-345 and the S339-341A mutation were encoded within the 5' ends of primers MJC261 and

MJC262 to provide a seamless introduction of the mutation in the PNPase 1-713 ORF in the Gibson reaction. Primers MJC222 and MJC223 include overhang sequences homologous to regions flanking the NheI site in pTEV5. Gibson assembly reaction was completed as previously described resulting in plasmid pMJC0112 which incorporates S339-341A into PNPase. Plasmid pMJC0113 was generated similarly with a modification to the second PCR fragment (PNPase codons 337-713), which was amplified from pMJC0095 with MJC262 and MJC226 to include the C-terminal flexible linker (LRSRAPENSNVTRHRSAT) and mCherry.

**Plasmid pMJC0119, RNase CTD PNPase binding site mutant expression vector:** pMJC0119 encoding RNase E-CTD-ΔPNPase_BS-eYFP contains the C-terminal domain (residues 451-884) of RNase E with a deletion of the PNPase binding site (residues 885-898) and a C-terminal linker (LRSRAPENSNVTRHRSAT) followed by eYFP was generated in pTEV5 using the Gibson assembly method with primers designed by j5 DNA assembly design software. PCR amplified RNase E codons 451-884 from pDT177 with primers MJC265 and MJC280. The linker region and eYFP were amplified by PCR from pDT177 with primers MJC281 and MJC269. Primers MJC280 and MJC281 include overhang sequences homologous to residues 880-884 and flexible linker codons encoding LRSRAP, and primers MJC265 and MJC269 include overhang sequences homologous to regions flanking the NheI site in pTEV5. Gibson assembly was completed as previously described resulting in plasmid pMJC0119, which deletes the PNPase binding site residues from RNase E-CTD-eYFP.

**Plasmid pMJC0137, PNPase-V104A/E224A/F233A-mCherry expression vector:** pMJC0137 encoding PNPase-V104A/E224A/F233A-mCherry contains PNPase with alanine mutations to key residues in the RNase E binding pocket of PNPase. pMJC0137 was generated in pTEV5 using the Gibson assembly method with primers designed by j5 DNA assembly design software. Each mutation was successively generated by site directed mutagenesis using helper plasmids. First, plasmid pMJC0121 was made by inserting mutation F233A into PNPase-mCherry using primers MJC282 and MJC283. Second, plasmid pMJC0134 was made by inserting mutation V104A into PNPase-F233A-mCherry using primers MJC290 and MJC291. Thirdly, the final plasmid pMJC0137 was made by inserting mutation E224A into PNPase-V104A/F233A-mCherry using primers MJC292 and MJC293. Gibson assembly was completed as described previously.

**Plasmid pDT279, MBP-RNase CTD expression vector:** pDT279 encoding MBP-RNase E-CTD-eYFP contains an N-terminal MBP fused to the C-terminal domain (residues 451-898) of RNase E with a C-terminal linker (LRSRAPENSNVTRHRSAT) followed by eYFP was generated in pTEV5 using the TEDA method[4] with primers designed by j5 DNA assembly design software. PCR amplified MBP from pTEV6 (pKLD66)[1] with primers DT471 and DT473. RNase E CTD, linker region, and eYFP codons were amplified using primers DT476 and DT477. 20 µL reactions containing 100 mM Tris-Cl pH 7.4, 10 mM $MgCl_2$, 10 mM DTT, 5 wt% PEG(8000), and 0.04 U T5 exonuclease were incubated with a 1:2 vector:insert molar ratio and 100 ng vector.


(1) Rocco, C. J.; Dennison, K. L.; Klenchin, V. A.; Rayment, I.; Escalante-Semerena, J. C. Construction and use of new cloning vectors for the rapid isolation of recombinant proteins from Escherichia coli. *Plasmid* **2008**, *59* (3), 231-237. DOI: 10.1016/j.plasmid.2008.01.001.
(2) Gibson, D. G.; Young, L.; Chuang, R. Y.; Venter, J. C.; Hutchison, C. A., 3rd; Smith, H. O. Enzymatic assembly of DNA molecules up to several hundred kilobases. *Nat Methods* **2009**, *6* (5), 343-345. DOI: 10.1038/nmeth.1318.
(3) Chen, J.; Densmore, D.; Ham, T. S.; Keasling, J. D.; Hillson, N. J. DeviceEditor visual biological CAD canvas. *J Biol Eng* **2012**, *6* (1), 1. DOI: 10.1186/1754-1611-6-1. Ham, T. S.; Dmytriv, Z.; Plahar, H.; Chen, J.; Hillson, N. J.; Keasling, J. D. Design, implementation and practice of JBEI-ICE: an open source biological part registry platform and tools. *Nucleic Acids Res* **2012**, *40* (18), e141. DOI: 10.1093/nar/gks531. Hillson, N. J.; Rosengarten, R. D.; Keasling, J. D. j5 DNA assembly design automation software. *ACS Synth Biol* **2012**, *1* (1), 14-21. DOI: 10.1021/sb2000116.
(4) Xia, Y.; Li, K.; Li, J.; Wang, T.; Gu, L.; Xun, L. T5 exonuclease-dependent assembly offers a low-cost method for efficient cloning and site-directed mutagenesis. *Nucleic Acids Res* **2019**, *47* (3), e15. DOI: 10.1093/nar/gky1169.
